# Perceptions of Equity, Challenges, and Identity-Based Differences Among U.S. Entomologists

**DOI:** 10.64898/2026.08.18.745652

**Authors:** S. Solange Barros-Bustos, Charles I. Abramson, Ana M. Chicas-Mosier

## Abstract

This study aimed to examine entomologists’ perceptions of equity, inclusion, and exclusion within the discipline, identifying perceived challenges and proposed solutions for advancing equity in the field. The present study surveyed 47 self-identified entomologists living in the United States in 2023. Using a mixed-methods design, the study examined entomologists’ perceptions of inclusion and exclusion within the discipline through qualitative and quantitative measures. Respondents highlighted needs for race-conscious funding opportunities, comprehensive inclusion efforts through geographically diverse outreach, and access to role models and mentors with similar identities to future entomologists. Findings are discussed in relation to a smaller 2013 study on recruitment and retention of entomologists of color, which offers preliminary historical context. The 2013 respondents emphasized intrinsic and age-based barriers to recruitment (e.g., lack of interest, limited K–12 outreach), the 2023 findings point toward structural inequities and retention needs (e.g., systemic exclusion). The 2013 comparison is interpreted as exploratory given differences in sample size and scope between the two studies. The 2023 study highlights evolving perceptions of persistent inequities in entomology and identifies opportunities to build a more inclusive and representative discipline.

## Introduction

Gender and racial representation disparities remain imbalanced in Science, Technology, Engineering and Mathematics (STEM) fields (1). Exclusion in these disciplines in the U.S has deep historical roots, with Black, Indigenous, and people of color (BIPOC) being systematically marginalized since the earliest days of American higher education (2,3). This continued structural and systematic underrepresentation can be described as a “leaky pipeline,” in which marginalized students leave STEM fields at disproportionately high rates throughout their education and careers (1).

Black and Latina/o/x students are ∼19% and 13%, respectively, more likely than White students to leave STEM majors for non-STEM fields (4). This disparity persists even after accounting for social background and institutional factors. Among students who initially declared STEM majors, only 34% of Black and 43% of Latina/o/x students complete a bachelor’s STEM degree, compared to 58% of White students (4). Broader attainment gaps also persist, with Black, Latina/o/x, and Indigenous students earning STEM bachelor’s degrees at rates lower than their relative representation in the U.S. population (5).

Several interconnected factors contribute to these disparities. Systemic barriers begin in K–12 education environments including disparities in teacher quality, school funding, and access to advanced math and science courses, leaving many students from historically excluded and underrepresented (HEU) backgrounds less prepared for rigorous college-level STEM work (6). In university, HEU students often report feelings of alienation in STEM due to a lack of representation in faculty, classrooms, and educational materials (3). Moreover, STEM cultures are frequently described as having a “chilly climate,” where prevailing White, masculine norms can feel unwelcoming or discriminative for women and HEU students (7).

Efforts to address inequities in STEM are shaped by differing views on the role of race in educational and professional outcomes. One perspective described in the literature is color-evasiveness, defined as the belief that ignoring race will reduce racism. Scholars have argued that color-evasive approaches can obscure structural inequities and discourage race-conscious efforts to address disparities (8,9).

A 2011 National Academies report emphasized the importance of increasing participation of groups underrepresented in the STEM workforce, both to expand economic opportunity across the U.S. population and to address the growing demand for STEM-trained professionals (10). A subsequent 2019 report, reinforced that “diversity in the workplace both expands the available talent pool and increases the range of perspectives and expertise available to solve grand challenges in STEM” (11), p. 31). Despite these calls, severe inequities persist.

Entomology, situated at the intersection of ecology, evolutionary biology, and agriculture, reflects many of the disparities seen across STEM fields. The Entomological Society of America (ESA), frames diversity, equity, and inclusion (DEI) as “a call-to-action that requires short-term improvements and long-term strategic planning…” In July 2015, ESA’s Governing Board established a Diversity and Inclusion (D&I) committee to propose resources, programs, and services that would maximize diversity and inclusion among ESA’s population and increase members’ cultural competency (12). In 2024, ESA membership demographics were predominantly White, with 60.9% of members identifying as White/Caucasian/European American. Asian/Asian American (10.9%) and Hispanic/Latino/a (8.3%) while all other racial and ethnic identities each accounted for less than 3% of membership (12).

These membership patterns mirror broader disparities across the U.S. science and engineering (S&E) education pipeline, where representation of historically excluded groups tends to decline at higher degree levels. In 2019, Black/African American, Hispanic/Latino/a individuals were underrepresented in their share of science and engineering bachelor’s degree earners compared to their share of the U.S. population. In contrast, White and Asian individuals were overrepresented in the same metrics (13).

In 2019, doctorates earned by Black/African American individuals were most highly concentrated in medical and other health sciences (39.7%), followed by social sciences and psychology (31.6%). In contrast, biological and agricultural sciences, in which entomology is most closely situated, accounted for only 11.9% of doctorates earned by Black individuals. A notable pattern is that biological and agricultural sciences ranked as the second most common doctoral disciplines among American Indian or Alaska Native, Hispanic/Latino/a, and Asian individuals (Table 1)(13).

**Table 1.** Percent of S&E doctoral degrees earned, by field, citizenship, and selected race or ethnicity in 2019.

| **Race, ethnicity, or citizenship** | **Engineering, Computer Science, Math, and Statistics** | **Biological and Agricultural Sciences** | **Medical & Health Sciences** | **Physical, Earth, Atmospheric, & Ocean Science** | **Social Sciences and Psychology** |
| --- | --- | --- | --- | --- | --- |
| **American Indian or Alaska Native** | 17.1% | 26.1% | 20.7% | 8.1% | 27.9% |
| **Black or African American** | 12.3% | 11.9% | 39.7% | 4.5% | 31.6% |
| **Hispanic or Latino\*** | 17.1% | 24% | 15.1% | 10.1% | 33.7% |
| **White** | 21.1% | 22.6% | 19.1% | 13.9% | 23.4% |
| **Asian** | 32% | 24.6% | 17.1% | 10.8% | 15.5% |
| **Temporary visa holders** | 53.7% | 14.9% | 4.8% | 14.8% | 11.8% |

Notes: Adapted from S&E doctoral degrees earned, by field, citizenship, and selected race or ethnicity: 2019, by National Science Foundation & National Center for Science and Engineering Statistics, 2019.
\*Hispanic may be of any race; race categories exclude Hispanic origin. Race or ethnicity categories apply only to U.S. citizens and permanent residents.

Likely, many of the barriers contributing to exclusion and self-selection out of non-medical biological science also drive underrepresentation in entomology and are further compounded by challenges common in the agricultural sciences. O’Brien et al. (6) found that Black/African American students had less exposure to ecology, reported fewer same-race role models, felt less comfortable in outdoor environments, were more religious than White students, and had more communal goals. These factors may pose barriers by limiting early exposure to ecology, reducing comfort in research environments, and constraining access to role models, while also contributing to cultural mismatches which weaken students’ sense of belonging in the discipline (6). However, recent work found no evidence that historical associations between agriculture and racialized labor negatively influenced contemporary student perceptions of entomology or agricultural sciences (14).

A recent study examined undergraduate perceptions across nine academic disciplines, including entomology, using measures of difficulty, safety, importance, relevance, beauty, and job availability. Across the full sample, entomology was generally perceived less favorably than other STEM disciplines, although it was also viewed as relatively welcoming, nonintimidating, and less difficult than fields such as physics or genetics (14). When non-White respondents were pooled into a single ‘people of color’ (POC) category, significant differences emerged in fear-related perceptions of biology, genetics, and entomology. Respondents in the POC group reported higher fear scores (Likert-scale range: 0= “no danger”, 3= “very dangerous”) for biology, genetics, and entomology than White respondents (14). The study also identified racial variation in career motivations and interests. Black respondents were more likely to report altruism and social improvement as career motivations, expressed interest in working with people, and rarely indicated interest in working with animals (<4%). Notably, no Black respondents reported an interest in working with insects while all other racial groups showed some level of interest in working with insects. By contrast, White respondents were primarily motivated by altruistic or social improvement and financial access.

These findings highlight the need to examine why entomology and agricultural science are perceived negatively by students and to identify leverage points for improving recruitment and retention in these disciplines. Previous studies have suggested strategic and geographically diverse outreach, framing entomology as an altruistic discipline, and promoting access to and retention of same-race mentors and faculty have been suggested by various studies (15); (16). However, participants have not been explicitly asked about challenges to recruitment and retention in the discipline since 2013.

Faculty mentoring can be positively associated with student satisfaction regardless of the mentor’s ethnicity (17,18). Despite its importance, mentorship often lacks formal recognition within higher education; for example, only 22% of surveyed science and engineering majors strongly agree that they had a mentor during their undergraduate studies (19). Less than 10% of faculty in STEM-related departments in U.S. public universities are people of color. Within the biological sciences, representation is even lower with <5% faculty of color. Such discrepancies in racial representation limit access to same-race mentors and role models in biology-adjacent training pathways, including entomology, contributing to persistent challenges in recruitment, retention, and recognition of BIPOC scientists (20).

### Gender in Entomology

Gender parity in biology-related disciplines has increased considerably over recent decades. In 2019, women earned 51.9% of doctoral degrees in the biological sciences and 49.7% in agricultural sciences, both higher than the overall average for all science and engineering fields (45.8%) (13). Gender parity does not necessarily equate to gender equity; systemic biases continue to limit STEM gender representation (21). Grogan (2019) showed that women in entomology remain underrepresented in employment relative to their representation among doctoral degree recipients. Although women earned at least 40% of U.S. entomology doctorates over the previous decade, they held roughly 25% of entomology positions in academia and the federal government in 2015–2016. Men are also disproportionately represented in higher-ranking positions, whereas women are concentrated in lower-ranking positions. Within the federal government, women entomologists received lower mean salaries at the highest grade level. At the current pace, gender parity is not expected to be achieved in the academy for many decades (23). While the traditional “leaky pipeline” framework emphasizes attrition at sequential career stages, more recent work highlights how gender inequalities are embedded across critical STEM processes (20,24). Factors that shape career progression even after entering academia such as, funding, hiring, and recognition are still challenged by systemic inequities Perceptions of entomology may be less impacted by respondent gender than other STEM disciplines. Evangelista et al. (14) found that gender was associated with respondents’ perceptions of academic fields but, these differences were minimal for entomology-related areas. Agricultural science showed gendered differences in perceived fear, with gender non-conforming students reporting more fear than males. There were no gender or sexual identity differences in the perception of entomology or ecology. However, LGBTQIA+ respondents rated entomology as more “beautiful” than heterosexual respondents but less approachable (14).

### Historical Context for this Manuscript

A 2013 study explored perspectives of Black and Hispanic Entomologists on STEM recruitment and retention (25). That study marked a significant step in documenting the experiences, challenges, and perspectives of POC within the field of entomology and brought attention to systemic barriers and the urgent need for inclusive recruitment strategies in STEM, particularly for African American/Black and Hispanic scientists. The researchers found a notable paradox: while most respondents strongly supported efforts to recruit Black and Hispanic students into entomology, they were less convinced that ethnicity itself should be a deciding factor in recruitment strategies. Respondents supported integration of entomology into high school curricula, setting up exhibits in inner-city libraries, and the creation of an interactive website designed by Black and Hispanic entomologists as effective ways to boost recruitment and that early tutoring in math and science could help increase representation. Positive mentor relationships emerged as the most referenced factor to encouraging students to pursue entomology, with both groups supporting visiting mentor programs and more international collaboration led by Black and Hispanic entomologists. Participation in professional meetings and research presentations was also considered important. Although both groups believed government funding should support recruitment efforts, they showed weaker agreement on whether universities should provide identity-based scholarships or whether a national entomology curriculum was necessary (25).

All respondents emphasized the importance of early exposure to science, particularly through role models and mentors who share their cultural background. They noted that students often lack awareness of entomology as a career option and suggested that outreach efforts should begin in elementary or middle school. Many respondents described entering the field by chance rather than through structured recruitment, indicating a need for more deliberate pathways into the discipline. Barriers such as financial constraints, limited access to quality education, and underrepresentation in higher education were commonly mentioned. Several participants expressed a strong desire to give back to their communities by mentoring younger students and increasing visibility of entomologists of color. Others shared personal experiences of isolation or marginalization in academic or professional settings, reinforcing the need for supportive networks and inclusive environments within the field (25).

The present study builds on that foundational work, revisiting many of the same themes while investigating a new group of respondents and modernizing the methodology through a more inclusive lens. The present authors duly recognize that the previous work had challenges to its methodology, specifically estimating POC based on website imagery. Today, data are more readily available to estimate representation in the field of entomology and online communication allows for broader recruitment efforts. Using the 2013 study as historical context, the present study examines entomologists’ current perceptions of inclusion and exclusion in 2023, exploring whether previously identified trends appear to have persisted, how representation in entomology has evolved in the United States, and what new challenges or opportunities have emerged for people of color in the field. Given differences in sample size and scope between the two studies, comparisons with the 2013 findings are treated as exploratory and interpretive rather than as a formal longitudinal comparison.

### Research questions

1. What are perceived challenges and prospective solutions for increasing equity in entomology?
2. How do perception of inequities in entomology vary by individual identities?
3. In what ways do perceptions of (in)equities in entomology in 2023 appear to differ from those reported in the 2013 study?

## Method

This study employed a mixed methods design, combining both quantitative and qualitative survey data to explore the experiences and perspectives of historically excluded individuals in entomology. The study was approved by the University of Kansas Institutional Review Board (Study #: 00149311) prior to data collection.

### Participants and Sampling

Participants were recruited using snowball sampling, a non-probability technique particularly useful for reaching small or hard-to-access populations. Recruitment took place between January 4, 2023, and May 12, 2023. Initial participants were identified through professional networks and were encouraged to forward the survey link to colleagues who met the inclusion criteria. Eligibility was limited to individuals who were at least 18 years old, self-identified as an entomologist, and resided in the United States. A total of 57 individuals initially participated in the study. However, 10 were excluded from the final sample due to incomplete responses or early withdrawal from the study. Given the anonymous nature of the survey, participants were provided with an Information Statement in lieu of a signed consent form. The Information Statement outlined the study’s purpose, procedures, risks, and participants’ rights. Participation in the survey constituted implied consent.

Of the 47 responses, the majority identified as female (51.1% of the sample) and the largest age group was 25–34 (44.7%) (Fig 1).

**Fig 1.**
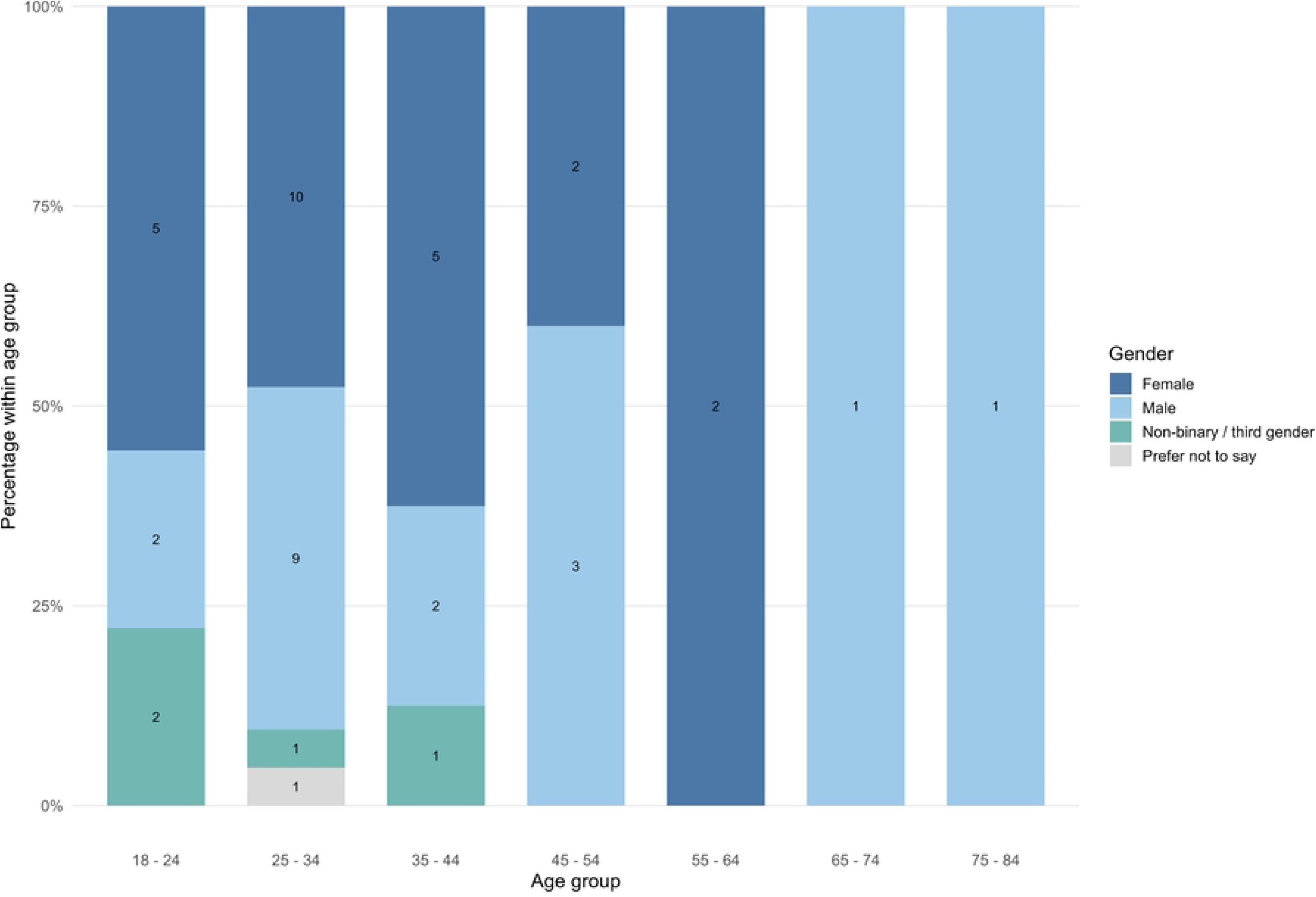
Age and gender distribution. Stacked bar charts display the demographic distribution of respondents by age and gender as percentages of the total sample.

### Survey Instrument

Data was collected using an online survey developed and administered through Qualtrics (2023). The instrument included both open- and closed-ended questions designed to explore participants’ attitudes, perceptions, and experiences related to inclusion, representation, and recruitment in the field of entomology. The survey included demographic questions, (race/ethnicity, gender, education level, and country of origin) followed by items related to participants’ professional experience, such as their current role, years in the field, and educational background.

The survey included 20 Likert-scale items rated on a 5-point scale ranging from 1 (*Strongly Disagree*) to 5 (*Strongly Agree*), along with eight open-ended questions. The Likert-scale items were organized into six subscales representing key attitudinal and perceptual domains related to diversity and inclusion in entomology. Six open-ended questions invited participants to elaborate on their experiences and offer suggestions for improving the recruitment and retention of scientists of color in entomology. Two items were reverse coded prior to analysis to ensure consistent interpretation of higher scores as reflecting more positive or inclusive attitudes.

***Experienced Workplace Discrimination*** (2 items): assessed the extent to which individuals perceived that they had personally encountered discriminatory treatment in professional settings based on race/ethnicity or gender.

***Positive Attitudes toward Inclusion*** (2 items): measured the degree to which respondents endorsed the value of diversity and supported inclusive practices within the field, such as recognizing the unique perspectives of people of color and considering race/ethnicity in recruitment.

***Awareness of Role Models and Diversity Resources*** (3 items): captured participants’ familiarity with role models and existing diversity-focused resources in entomology (e.g., Charles Henry Turner, online platforms highlighting entomologists of color).

***Perceptions of Need for Early Education*** (6 items): reflected beliefs about the importance of introducing entomology-related content and support structures into K–12 education to promote future interest and recruitment.

***Perceptions of Need for Outreach Strategies & Education*** (4 items): evaluated beliefs about the effectiveness of outreach initiatives, such as conferences, international collaborations, professional meetings, and online resources, in attracting underrepresented students to the discipline.

***Perceptions of Need for Financial and Institutional Support*** (3 items): assessed perceived necessity of financial and structural support (e.g., scholarships, tuition waivers, federal funding) aimed at increasing participation among HEU individuals in entomology.

### Psychometric analysis methods and procedures

*Quantitative Analysis*: Descriptive statistics were calculated to summarize the demographic characteristics of the sample, including race/ethnicity, educational attainment, professional affiliation, publication history, and career intentions. To assess participants’ perceptions and attitudes toward inclusion in entomology, six thematic subscales were computed from questionnaire items using 5-point Likert-type response options (1 = *Strongly Disagree* to 5 = *Strongly Agree*). Subscale scores were obtained by averaging the items within each theme.

To examine group differences, one-way analyses of variance (ANOVA) were conducted across race/ethnicity, gender, educational level, age group, and employment type for each of the six subscales. For the race/ethnicity variable, due to small sample sizes across most subgroups, categories other than Black or African American and White were combined into a single “Other” category for inferential analyses, resulting in three racial/ethnic groups: Black or African American (n = 9), White (n = 29), and Other (n = 9). This approach ensured adequate group sizes for one-way ANOVA and post-hoc comparisons.

Effect sizes were reported using eta squared (*η²).* When omnibus tests were statistically significant, Bonferroni post-hoc comparisons were applied to identify pairwise group differences. All statistical analyses were performed in SPSS 30 (27) unless otherwise specified.

*Qualitative Analysis*: We used a qualitative content analysis approach with quantitized frequency description to characterize the relative prominence of themes. Survey responses were exported from Qualtrics, compiled into a single dataset, and reviewed for completeness prior to analysis. Responses were de-identified by removing any potentially identifying information before coding. Each qualitative question was analyzed independently.

Two of the three authors independently coded the responses; both have backgrounds in educational research and STEM equity. Codes were developed iteratively through repeated reading of the responses using the Naeem et al. (2023) 6Rs framework (Robust, Reflective, Resplendent, Relevant, Radical, Righteous). Coding discrepancies were discussed until consensus was reached, and the final coding framework was refined collaboratively. Similar codes were grouped into broader categories and themes, resulting in hierarchical coding structures that were subsequently represented as dendrograms. A manifest content analysis was first conducted on the open-ended responses to identify recurring ideas and patterns explicitly stated by respondents. The process included: (1) familiarization with the data; (2) identification of meaningful keywords; (3) coding of data segments using the 6Rs framework (Robust, Reflective, Resplendent, Relevant, Radical, Righteous); (4) development of themes through grouping related codes; and (5) conceptualization of patterns into defined concepts. The sixth step, construction of a conceptual model, was not performed because the purpose of this analysis was to describe and interpret emergent patterns rather than to develop a formal model.

To assess the prominence of ideas, we assigned weights to each code based on frequency of occurrence across responses. After coding each response, the number of times a code appeared was aggregated across the dataset. Each occurrence was treated as one unit of weight, such that codes with higher frequencies received proportionally greater weights. These weighted frequencies allowed us to calculate the relative prominence of each code, expressed as a percentage of the total number of coded segments. Based on the code theme, the codes were grouped into broader categories and subcategories. Finally, we visualized these relationships using a dendrogram and summarized the distribution of codes with weighted percentages. This allowed us to identify which issues emerged most consistently in the dataset. Condensed dendrograms are presented in the main text to improve readability, whereas complete dendrograms containing all codes and hierarchical relationships are provided in the Supplementary Materials.

Two open-ended questions were analyzed comparatively: *What research areas do you feel should receive more attention in the next 20 years?* and *How has the paradigm of Entomology changed in the past 20 years?* These items were analyzed in R (29) using the packages *dplyr*, *readr*, *ggplot2*, and *tidyverse*. Frequency counts were calculated by grouping responses according to survey question and thematic category, resulting in a category-by-question summary table. To facilitate direct comparison between prospective priorities and retrospective paradigms, counts for the retrospective question were assigned negative values, allowing the thematic emphasis of the two questions to be contrasted on a common scale. The aggregated data were used to generate comparative visualizations, including side-by-side and diverging bar charts.

## Results

### Descriptive statistics

Of the 47 respondents, 61.7% identified as White and 19.1% as Black or African American. Smaller proportions identified as Asian (4.3%) and Hispanic/Latino/a (4.3%). For analysis, several identities were grouped into an “Other” category (10.6%), which included individuals who identified as American Indian or Alaska Native, half Mexican and half Polish, White and Black or African American, and White and Native Hawaiian or Pacific Islander (Fig 2).

**Fig 2.**
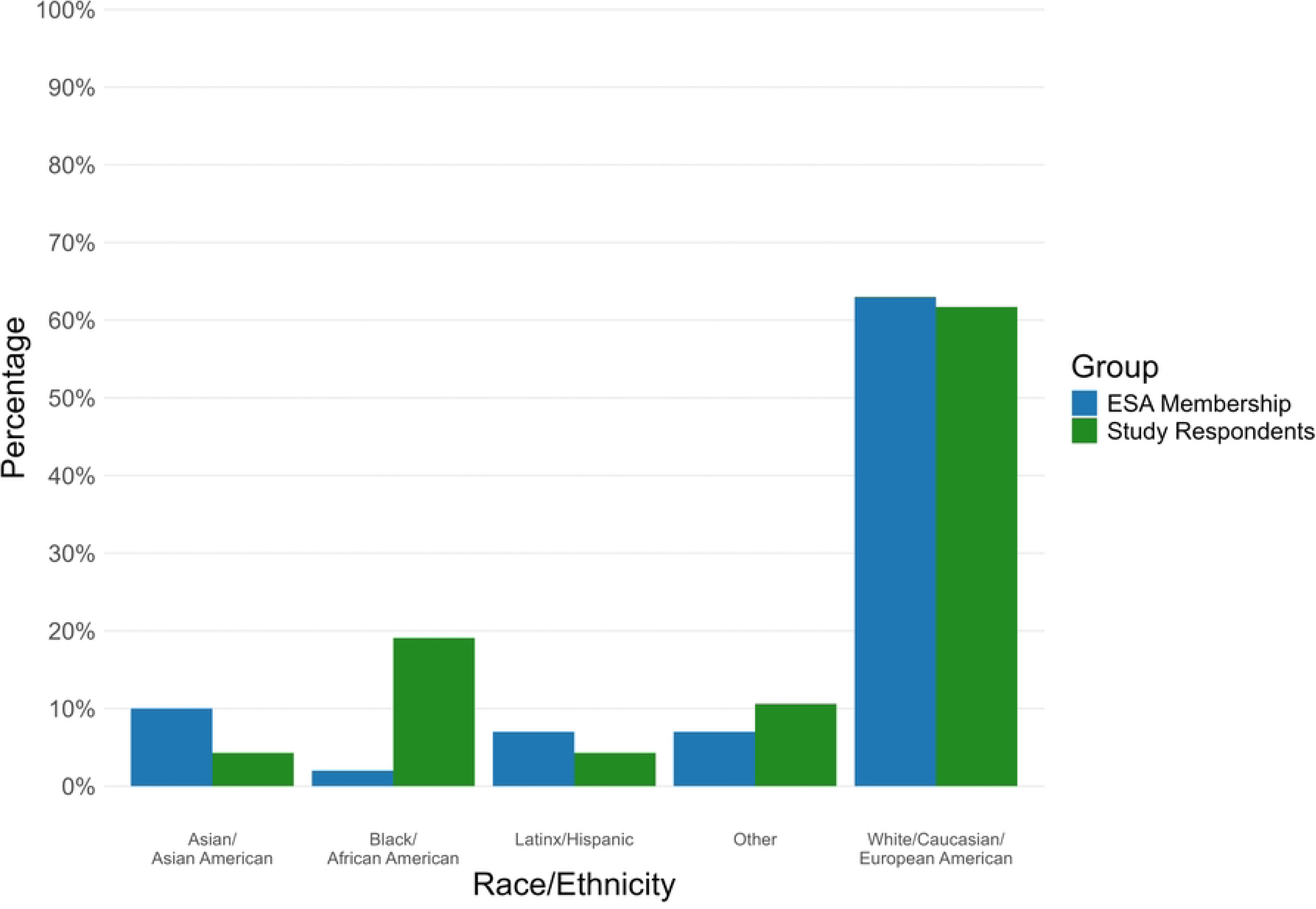
Race/ethnicity distribution. Race/ethnicity distribution of the present work compared to ESA membership (ESA, 2025)

Membership for the Entomological Society of America is 63% White; a nearly identical proportion was observed in this sample (Fig 2). However, Black or African American individuals made up 19.1% of this sample, compared to only 2% of ESA membership (ESA, 2025). Hispanic/Latino/a (4.3%) and Asian (4.3%) participants in this sample were slightly underrepresented relative to ESA’s reported 7% and 10%, respectively. Additionally, 10.6% of respondents in this study were categorized as “Other”. In comparison, ESA reports a combined 7% of members within similar “Other” categories, including African, Multiracial/Biracial, American Indian/Alaska Native/Indigenous/First Nations, Arab/Middle Eastern, Native Hawaiian/Pacific Islander, and Other. The present sample may offer a broader representation of racially and ethnically diverse voices, especially of Black respondents. Targeted oversampling such as this has been suggested as a means to promote racially representative sampling (30).

The sample was highly educated and largely academic (Fig 3). The largest group held doctoral degrees (38.3%), and most respondents were students (44.7%) or professors, staff, and administrators (36.2%) (Fig 4). Nearly half of those without a terminal degree intended to pursue one (44.7%), and most participants had at least one publication (74.5%).

**Fig 3.**
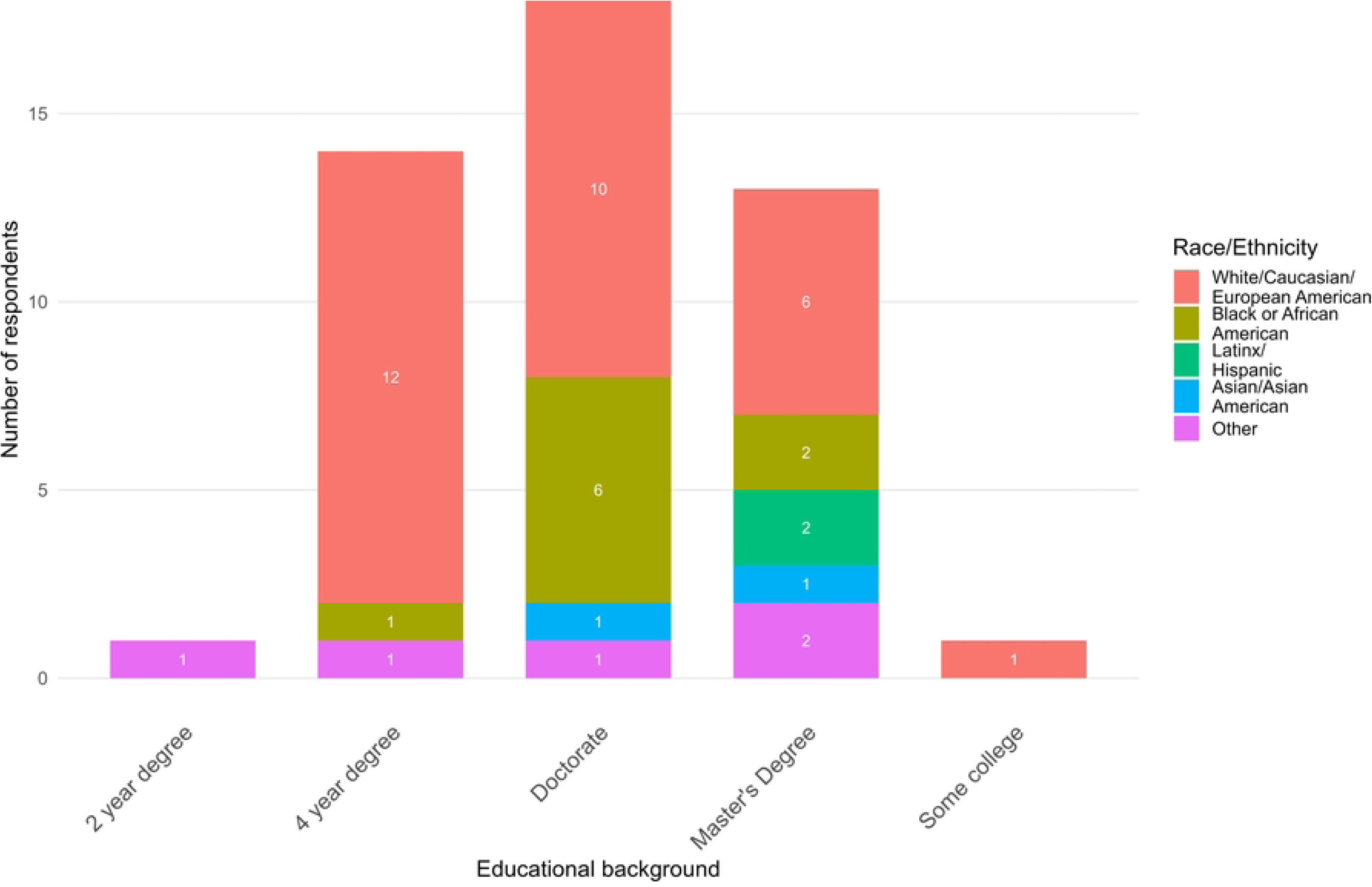
Educational background of survey respondents by race/ethnicity. Stacked bar chart showing the distribution of respondents across educational backgrounds, stratified by self-identified race/ethnicity. Numbers within each bar segment indicate the number of respondents in each category.

**Fig 4.**
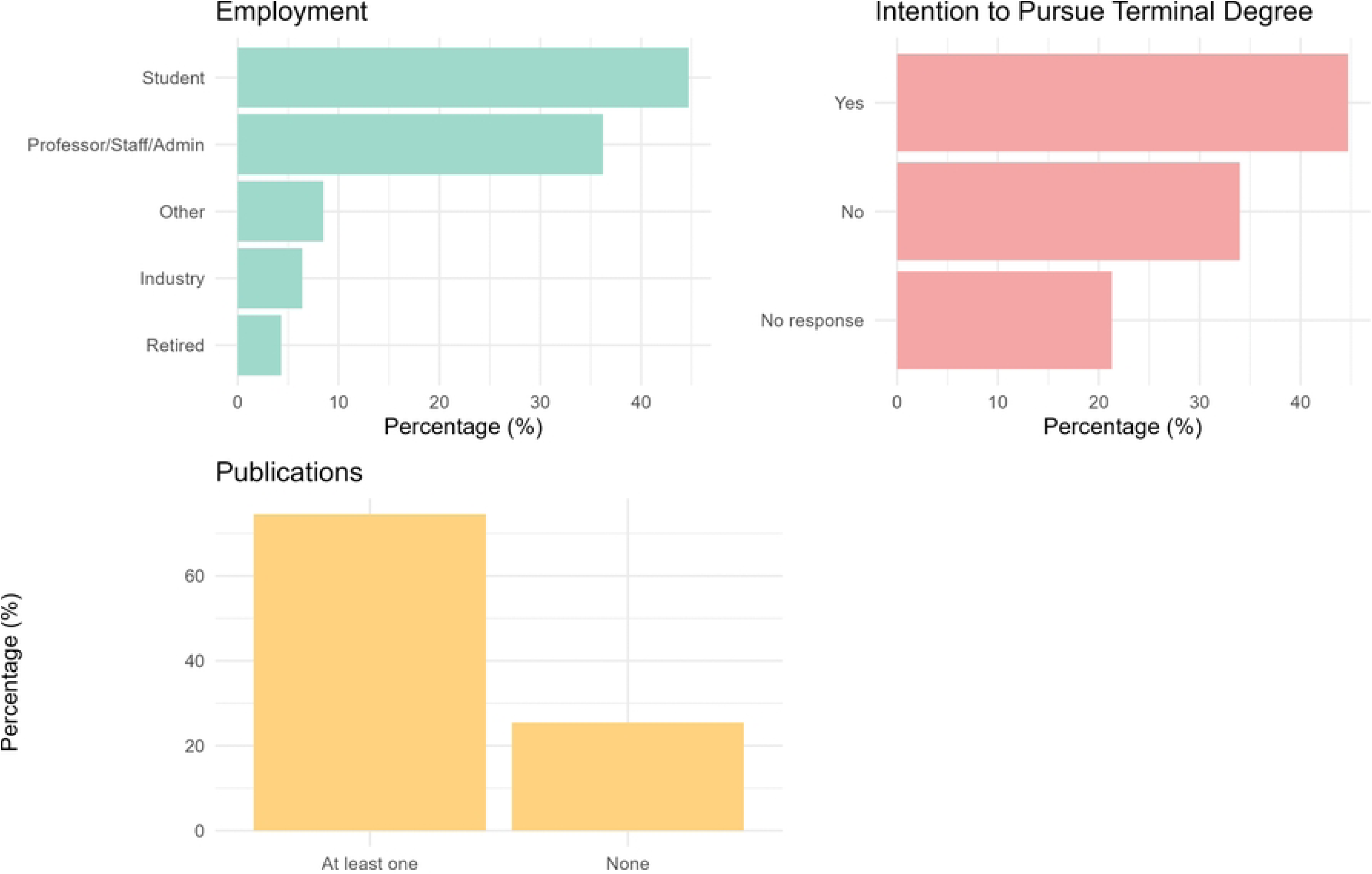
Employment status, intention to pursue a terminal degree, and publication experience among survey respondents. Distribution of respondents by employment status (top left), intention to pursue a terminal degree (top right), and publication experience (bottom). Values represent the percentage of respondents in each category.

Respondents were distributed across multiple regions of the United States. Based on self-reported birthplace (Fig 5), most respondents were born in the United States, particularly in the Midwest and Northeast. In addition, a smaller subset of participants reported being born outside the United States, including in South and Central America, West, Central, and South Africa, and Southern Asia, reflecting some degree of international origin within the sample. In addition, Qualtrics-generated geolocation data (latitude and longitude) reflected participants’ locations at the time of survey submission. Response-location data (Fig 6) indicate that respondents were distributed across the United States. Although participants were dispersed nationally, clusters of responses appeared in the Midwest, Southeast, and Northeast.

**Fig 5.**
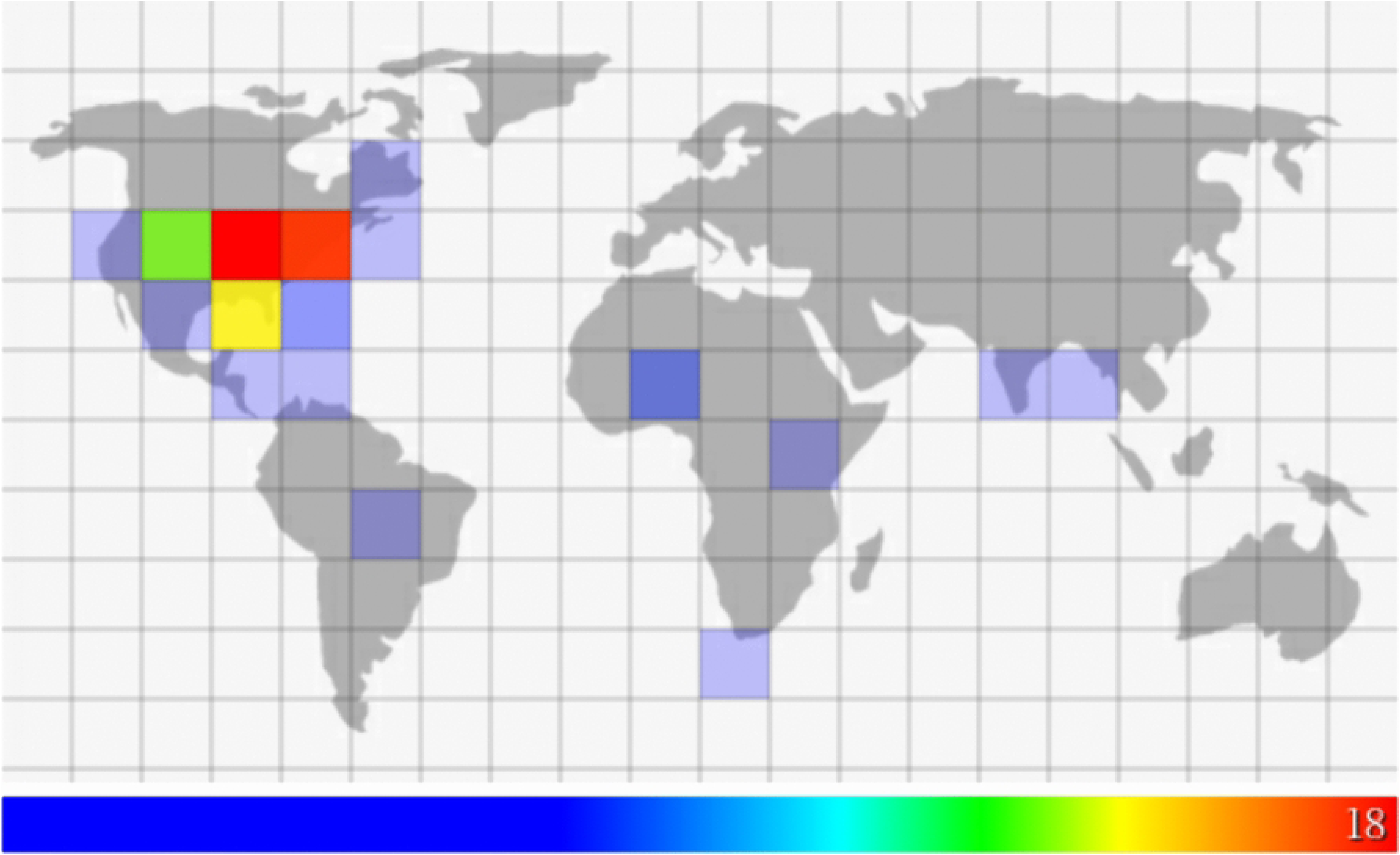
Self-reported birthplace of survey respondents. Geographic locations reflect respondents’ answers to the question: Where were you born (click the area on the map)? Colors indicate the frequency of responses within each geographic region, with warmer colors representing higher concentrations (max=18 respondents).

**Fig 6.**
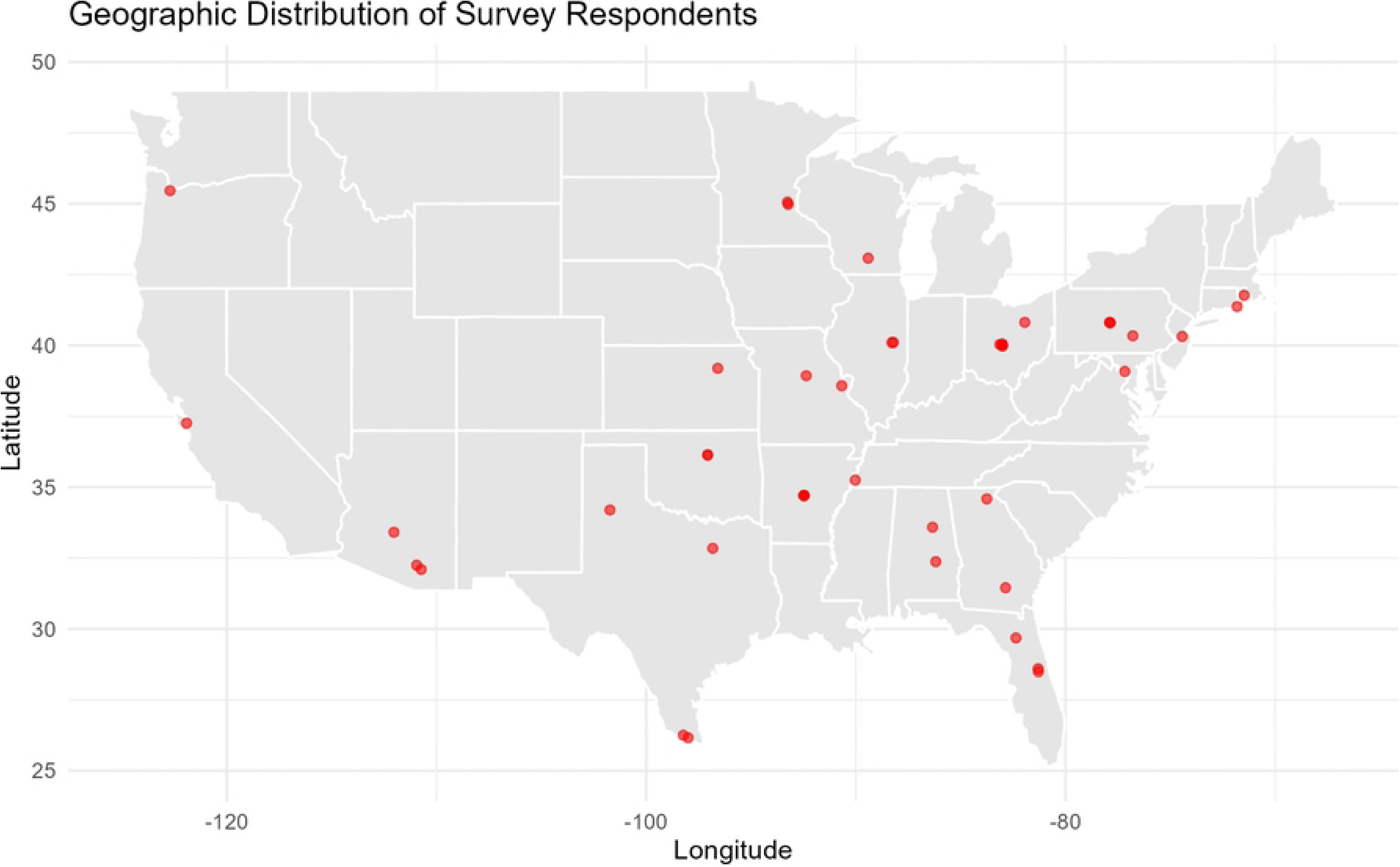
Geographic distribution of survey respondents at time of survey submission. Locations are based on latitude and longitude coordinates automatically recorded by Qualtrics at the time of survey completion. Points represent respondents’ approximate current location within the United States.

### Quantitative Analysis

Table 2 presents the means and standard deviations for the six thematic scales assessing perceptions and attitudes toward inclusion in entomology. The highest mean value emerged in the Perceptions of Financial and Institutional Support theme (*M* = 4.19), reflecting strong endorsement of financial support mechanisms for inclusion, such as targeted funding and scholarships. The Attitudes toward Inclusion theme also scored highly (*M* = 3.92).

**Table 2.** Descriptive statistics for perception and attitude subscales related to inclusion in entomology.

| Theme | Mean | Std. Deviation |
| --- | --- | --- |
| Awareness of Role Models | 2.447 | 1.246 |
| Experienced Workplace Discrimination | 2.650 | 1.334 |
| Perceptions of Need for Early Education | 3.763 | 0.448 |
| Positive Attitudes toward Inclusion | 3.915 | 0.974 |
| Perceptions of Need for Outreach Strategies & Education | 3.946 | 0.662 |
| Perceptions of Need for Financial and Institutional Support. | 4.185 | 1.006 |

Notes: Mean and Standard Deviation for the six perception and attitude subscales. Higher scores indicate greater endorsement of each construct.

In contrast, the lowest mean was observed in the Awareness of Role Models (*M* = 2.45, *SD* = 1.25), indicating limited familiarity with figures such as Charles Henry Turner or with diversity-focused platforms like Entomologists of Color. This was followed closely by the theme Experienced Workplace Discrimination (*M*= 2.65, *SD*= 1.3). Supplementary Material lists the questions included in each subscale.

A one-way ANOVA was conducted to examine differences across ethnic groups (Black or African American, White, and Other) on the six perception and attitude subscales related to inclusion in entomology (Table 3). Significant group differences emerged for Perceived Workplace Discrimination, *F*(2, 44) = 4.82, *p* =0.013, *η^2^* = 0.180, 95% CI [0.009,0 .350]. Bonferroni post-hoc comparisons indicated that Black or African American participants reported higher levels of perceived workplace discrimination *(M* = 3.78, *SD* = 1.28) than White (*M* = 2.45, *SD* = 1.17, *p* = 0.022) and Other respondents (*M* = 2.17, *SD* = 1.41*, p* = 0.025); White and Other did not differ significantly (*p* = 1.000).

Significant group differences were also observed for Awareness of Role Models, *F*(2, 44) = 4.93, *p* = 0.012, *η^2^* = 0.183, 95% *CI* [0.010, 0.353]. Black or African American respondents reported higher awareness (*M* = 3.48, *SD* = 1.48) than White (*M* = 2.30, *SD* = 0.94, p = 0.030) and Other respondents (*M* = 1.89*, SD* = 1.41, p = 0.016); White and Other did not differ significantly. For Perceptions of Need for Outreach Strategies & Education, the ANOVA revealed significant group differences, *F*(2, 43) = 3.56, *p* = 0.037, *η^2^* = 0.142, 95% *CI* [0.000, 0.310]. Black or African American participants reported the highest mean (*M* = 4.44, *SD* = 0.46), differing significantly from White respondents (*M* = 3.84, SD = 0.61, *p* = 0.044); the comparison with Other respondents did not reach significance (*p* = 0.106), nor did White versus Other (*p* = 1.000).

No significant group differences emerged for Attitudes to Inclusion, *F*(2, 44) = 1.09, *p* = 0.347, *η² =* 0.047; Perceptions of Need for Early Education, *F*(2, 43) = 1.30, *p* = 0.283, *η^2^* = 0.057; or Perceptions of Need for Financial and Institutional Support, *F*(2, 42) = 0.42, *p* = 0.657, *η^2^* = .020.

**Table 3.**
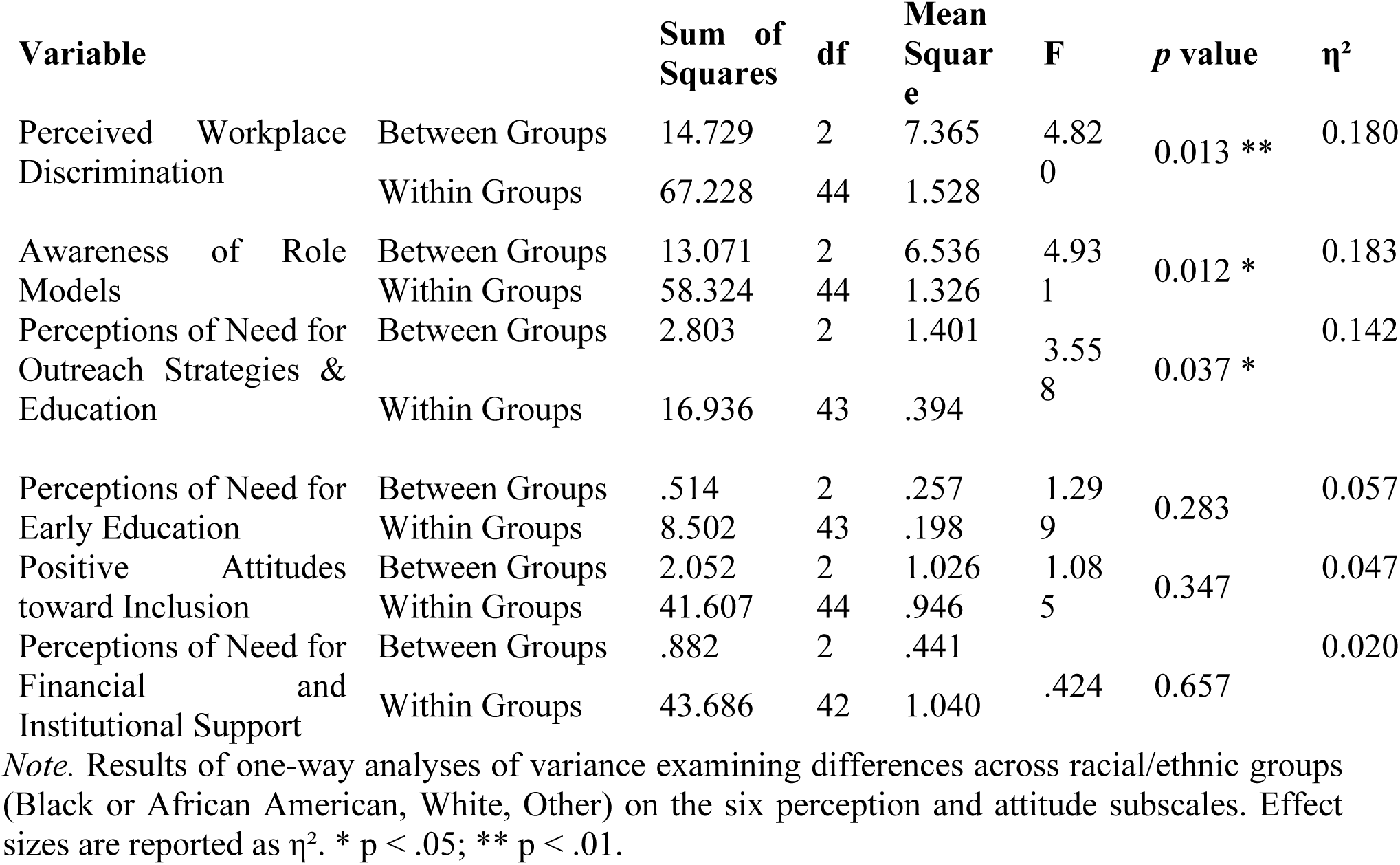
One-way ANOVA results for differences across ethnic groups (collapsed: Black.

A one-way ANOVA was conducted to compare the six themes of the scale by gender. The analysis revealed a significant difference in Perceived Workplace Discrimination (*F*(2, 44) = 4.048, *p* = 0.025, *η^2^* = 0.158). Bonferroni post-hoc comparisons showed differences between male (*M* = 2.03, SD=1.43) and female respondents (*M* = 3.15, SD= 1.20, *p* = 0.021). Notably, while the analysis included three gender groups, individuals identifying as Non-binary/Third Gender (*M* = 2.63, SD = 0.48) did not show significantly different levels of perceived discrimination compared to the other groups. Women reported experiencing the highest levels of workplace discrimination, highlighting an important gender-related disparity within the sample.

The analysis revealed a statistically significant effect on Awareness of Role Models by education level (*F*(4, 42) = 2.85, p = 0.035). Respondents with a doctorate degree reported the highest awareness (*M* = 3.15, *SD* = 1.34), followed by those with a master’s degree (*M* = 2.03, *SD* = 1.17) and a 4-year degree (M = 2.00, SD = 0.84). In contrast, those with a 2-year degree (*n* = 1*, M* = 1.33) and those with some college but no degree (*n* = 1, *M* = 2.67) showed less awareness. No significant differences were found across age groups or employment types in any of the six themes.

### Qualitative Analysis

#### What do you view as significant challenges for the future of Entomology?

The responses to this question were categorized into four broad areas of concern: Educational and Career Pathways (27%), Limited Outreach (19%), Scientific and Disciplinary Priorities (21%), and Diversity, Equity, and Inclusion (32%), (Fig 7)

**Fig 7.**
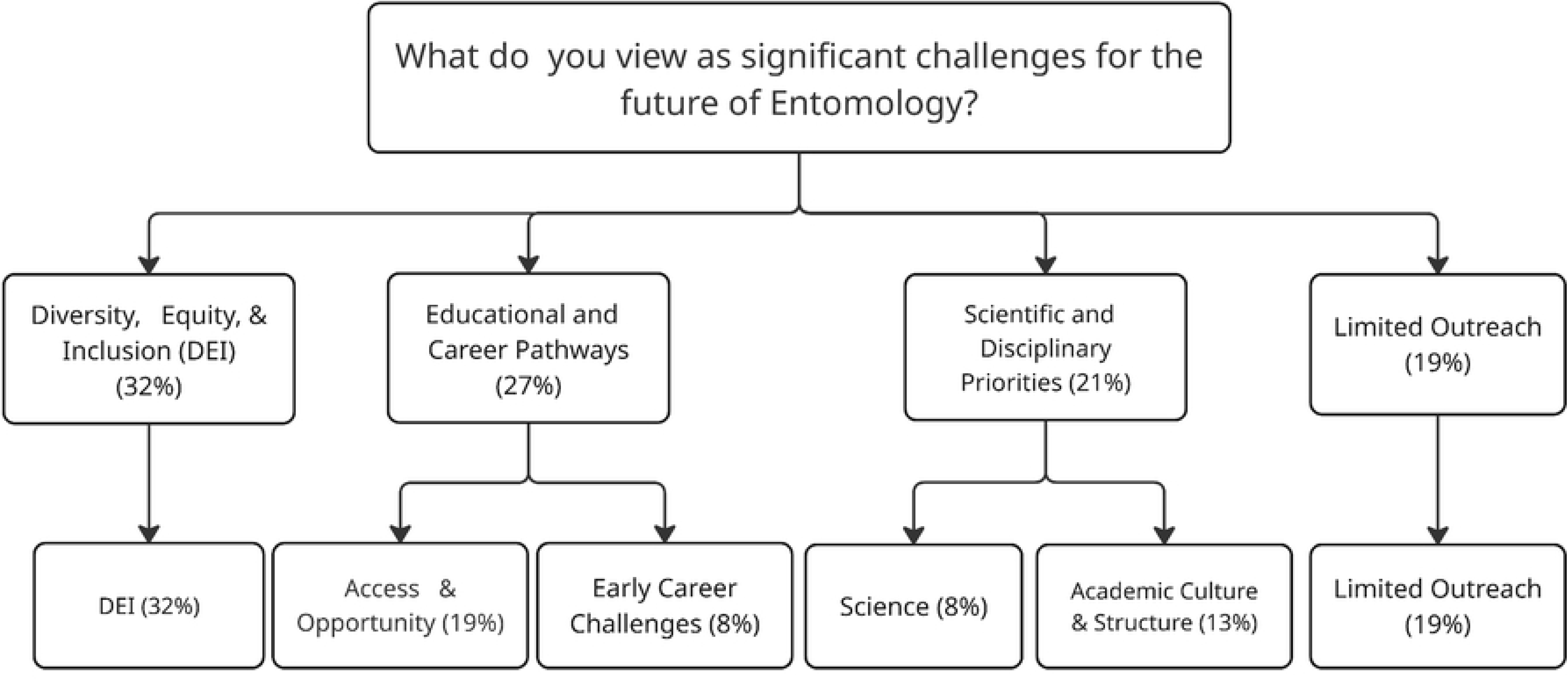
Thematic hierarchy of respondents’ perceived challenges for the future of entomology. Percentages represent the proportion of coded responses for each theme and subtheme.

Within Educational and Career Pathways, challenges such as funding (8%, e.g., “*Funding from the government”*), retention (4%, e.g., “*supporting students and young professionals so* [they] *enter the field and are retained”*), inadequate mentoring (3%), and mental health and wellbeing (3%, e.g., “*Graduate school is difficult on mental health*”) were recurrent themes, underscoring barriers to sustaining students and early-career professionals. Early Career Challenges such as lack of jobs (4%) and poor pay (4%) were also noted: “*I am seeing students being recruited, but not adequately mentored/paid or given an overload of responsibilities relative to their salary.*”

In Limited Outreach (19%), participants emphasized the importance of K–12 outreach (5%, e.g., “*Getting K-12 and high school students interested”*), recruitment (4%), and relatability (4%), along with addressing urbanization and technological distractions (3% each; e.g., “*Kids in suburban areas are not going outside as much as they used too* [sic]*, especially not without a screen”*) as factors that may limit future engagement with entomology.

Scientific and Disciplinary Priorities (21%) reflected concerns about both ecological challenges and academic structures. The “Science” subcategory (8%) captures core scientific challenges facing the discipline, particularly those related to environmental change and ecological pressures, such as climate change (3%), insect decline (4%), and pesticide impacts (1%), e.g., “*Decrease in insect abundance/diversity due to climate change*”. In the subcategory of Academic Culture and Structure (13%), the most cited issue was specializations within the field of entomology (8%) that reflected concerns about disciplinary narrowing and fragmentation, including pressures that may reduce breadth in training (e.g., “*fewer taxonomists or applied specialists”*). Respondents also linked these issues to training preparedness and scientific rigor, expressing concern that weak quantitative preparation among students could affect the quality of future entomological research. For example, “*students lack interest in learning math…”, “Need of high quality science”*. Respondents also offered structural critiques such as the disapproval of studying social topics in entomology (3%, e.g., “*Students have been wasting time studying social topics in science when these topics should be studied by students in sociology*”).

Finally, Diversity, Equity, and Inclusion (DEI) emerged as the largest area of concern (32%). Key challenges included racism (5%), unwelcoming environments (5%), and calls for diversity training (4%, e.g., “*Entomologists in positions of power need to have extensive diversity trainings*”). Additional barriers such as bias (3%), inclusion (3%), and access (4%) highlight the systemic issues facing marginalized groups in entomology.

#### What do you view as the most significant challenges in training young Black, Latinx, and Indigenous students?

In response to this question the sample identified a wide range of barriers, with the most frequently cited falling into five broad themes: Structural & Systemic Barriers (33%), Social & Cultural Exclusion (25%), Career Awareness & Exposure (32%), and Psychological & Motivational Factors (6%), (Fig 8).

**Fig 8.**
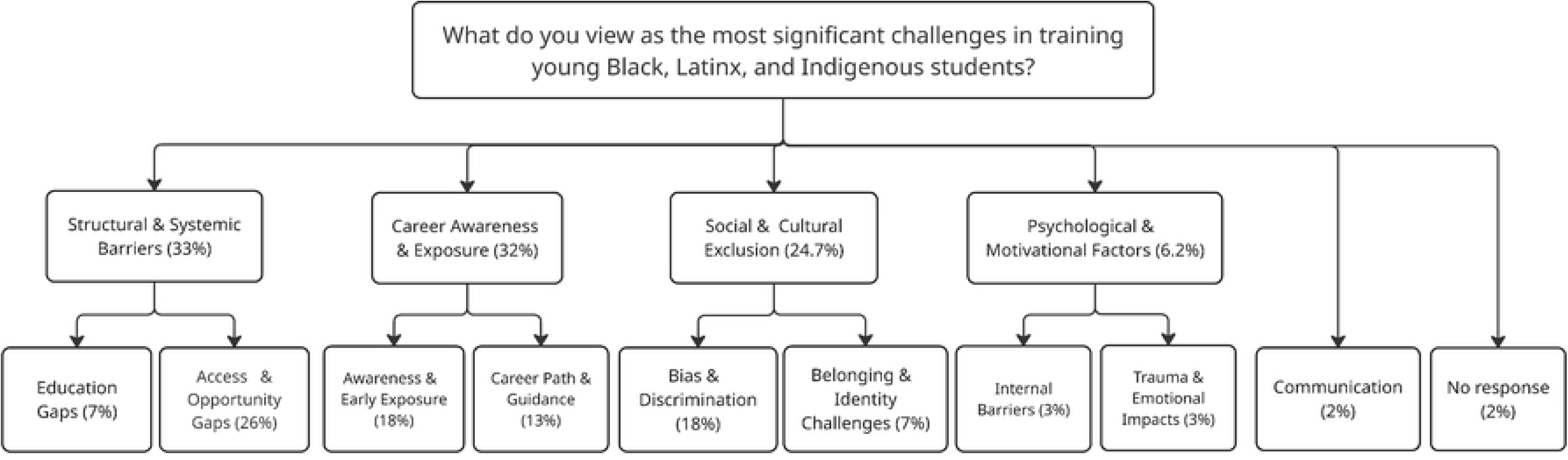
Codes and thematic categories for perceived challenges in training young Black, Latinx, and Indigenous students. Percentages represent the proportion of coded responses for each theme and subtheme.

In the Structural & Systemic Barriers category (33%) and subcategory of Access and opportunity Gaps (26%) respondents emphasized funding (8%, e.g., “*Money, which is the only way to disentangle the race issue with class/socioeconomic issues*”), financial barriers to access (4%, e.g., *Generational race related poverty puts individuals in the POC community at a significant disadvantage when entering higher academia, thwarting the ability for individuals to fund their own higher education*”), and broader resource gaps (2%) “*In low-income communities of color, more resources should be allocated to education in general and exposure to more job options, entomology included”,* as well as inequities in socioeconomics, access barriers, and the reliance on unpaid/volunteer opportunities. While in Education Gap (7%) they mentioned, Education Gaps (2%), First Gen Barriers (2%), and Class Barriers (2%) among others, with quotes like “*…POC are more likely to be first generation college students, and I have found this field is not welcoming to those first entering academia*.”

Social & Cultural Exclusion (24.7%) focused on bidirectional experiences of bias and discrimination, including color-evasive responses (5%, e.g., “*I have no idea, everyone I’ve met is super smart, I’ve had privilege of working with many different people*”), anti-racism (3%, e.g., “*Domestic students of color experience developmental trauma in the form of racism”*) and discussing access limitations from systemic exclusion such as sexism, transphobia, and institutional disrespect. Participants also highlighted a lack of belonging and representation (4%), noting that underrepresentation among faculty and mentors and a lack of culturally responsive support can make the discipline feel unwelcoming and reduce students’ sense of being represented (e.g., “*not having diverse teachers who understand cultural backgrounds*”).

Career Awareness & Exposure (32%) reflected two subcategories; Awareness and Early exposure (18%), which revealed perceptions that the most significant challenge in training is building interest (5%, e.g., “*Early exposure to entomology to remove social stigmas that are anti “bug*”) and entomology awareness (4%, e.g., “*Education, most people are not aware of the field*”). The absence of role models (1%) and negative perceptions of science (1%) may exacerbate barriers to entry.

Career Path & Guidance (13%) underscored the importance of mentorship support (4%, e.g., “*mentors who are genuinely engaged in recruitment*”) and clear career pathways (4%, e.g., *“Getting them exposed to the pathways to become an Entomologist*”), while also citing the need for additional professional development for career readiness and concerns of limited career options.

Finally, Psychological & Motivational Factors (6%) were mixed. While some rejected systemic challenges and criticized intrinsic motivations (2%, e.g., “*They* [HEU students] *start thinking that the society is in debt with them and that they are entitled to receiving more from society than other people*.”), others noted impacts like developmental (1%) or career trauma (1%, e.g., “*students of color experience developmental trauma in the form of racism; this trauma sets us back emotionally and developmentally*”). A smaller set of responses (2%) pointed to communication barriers, such as language or academic speech.

#### What strategies would you use to recruit Black, Latina/o/x, and Indigenous students into Entomology?

In response to this question, participants highlighted a wide range of approaches (Fig 9) including outreach (27%), Mentorship, Representation, and Support (17%), Cultural Support (23%), Financial Support (6%), Higher Education Strategies (18%), Barriers to Equity Thinking (6%), and Other (1%).

**Fig 9.**
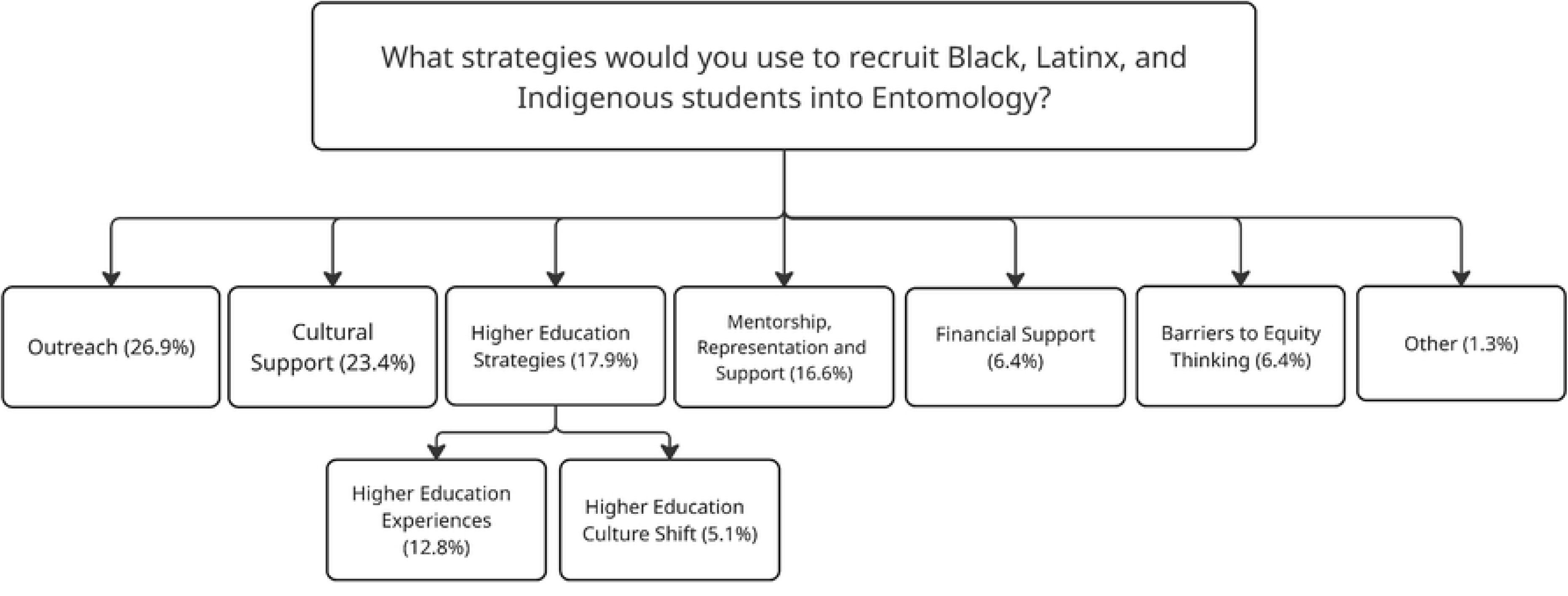
Dendrogram of responses to “What strategies would you use to recruit Black, Latina/o/x, and Indigenous students into entomology?” Percentages represent the proportion of coded responses for each theme and subtheme.

In the Outreach category, the most frequent mentioned strategy was K–12 outreach (26.9%), e.g., “*Doing outreach in more inner-city schools at all age levels, recruiting/ encouraging/ allowing interns from inner city high schools*”, underscoring the importance of early exposure to entomology as a pathway to recruitment.

Other recurring strategies included Mentorship, Representation, and Student Support (16.6%). In this category we can find creating role models (3.8%, e.g., “*including Black entomologists in learning standards (For example, Dr. Charles Henry Turner, Mrs. Sophie Lutterlough, Dr. Lonnie Standifer, Dr. Ernest J. Harris)*”, mentorship (5.1%, e.g., “*ensuring they have mentors or folks on their committee who are supportive*”) was seen as a critical mechanism to provide guidance and support.

In the category Higher Education Strategies (17.9%), responses were divided into strategies focused on institutional change and those centered on student experiences and opportunities. The Higher Education Culture Shift category (5.1%), captured strategies aimed at transforming institutional environments, including improving inclusivity (1.3%) and prioritizing Retention Over Recruitment (1.3%) *“I think it’s pointless to recruit BIPOC students if we are just going to suffer while in school*” and Strategic Job Posting (1.3%) *“Specific language included in job posting”.* In contrast, the Higher Education Experiences category (12.8%) reflected strategies that directly engage students through Research Opportunities (7.7%, “*Grade school and high school internship experiences*”), Expand Career Opportunities (2.5*%, “I would create paid positions for BIPOC students*”), and Education (2.5%).

Although less frequent, several respondents pointed to what we could consider “Barriers to Equity Thinking” (6.4%) where we can find codes like color evasiveness (2.5%), individuality and meritocracy (2.5%) and quotes such as “*Treat everyone with respect and as an individual and not a representative of a group. Do not patronize others*.” This quote was reiterated as a response to multiple questions by one respondent. Financial support (6.4%) included codes like “*I would highly recommend the increased funding towards college educational scholarships for POC in STEM fields*.”

#### Question 6: What, if any, barriers exist to recruitment of Black, Latina/o/x, and Indigenous students into Entomology?

The analysis of responses to the question “*What, if any, barriers exist to recruitment of Black, Latinx, and Indigenous students into Entomology?”* yielded 72 coded segments distributed across seven categories (Fig 10). The most frequently identified barriers were Economic and Labor Barriers (23.6%), followed closely by Representation and Mentorship Gaps (22.2%). Structural and Systemic Barriers (16.7%), Identity-Based Discrimination (13.9%), Outreach (12.5%), and Social Justice Avoidance (9.7%). A small proportion of responses (1.4%) reflected Field-Specific Perceptions.

**Fig 10.**
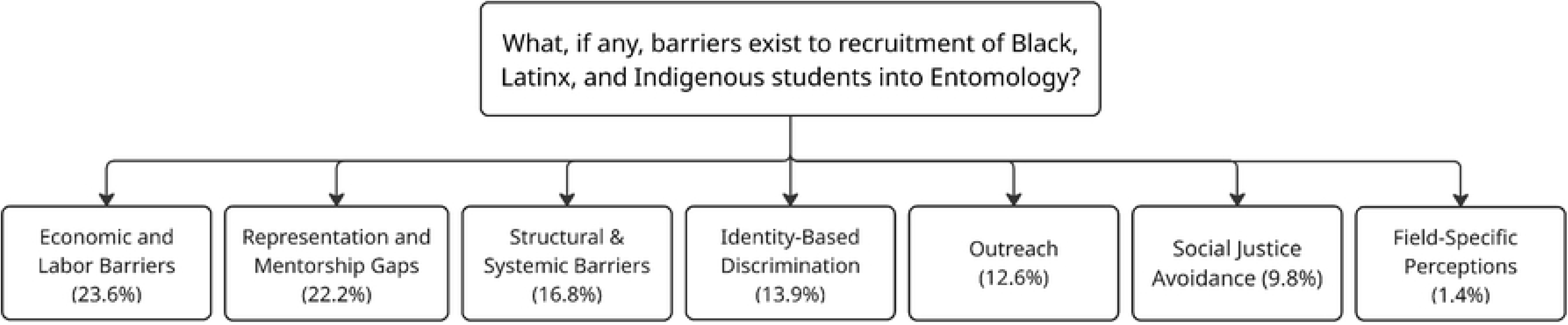
Dendrogram of responses to “What, if any, barriers exist to recruitment of Black, Latina/o/x, and Indigenous students into entomology?” Percentages represent the proportion of coded responses for each theme and subtheme.

Within Economic and Labor Barriers, funding was the most frequent individual code (11.1%), followed by social stigma (6.9%). Respondents referred to low compensation, unstable funding, and excessive workloads as structural deterrents to recruitment. As one participant stated, “*Low compensation vs. high work hours. It is difficult for grad students to have a second job to supplement our income*.” Others emphasized broader structural inequities, noting that “*Due to generational poverty and generational discrimination, POC individuals are much less likely to fund a college education*.”

Representation and mentorship gaps accounted for 22.2% of responses. The importance of role models (9.7%) was referenced with quotes like “*There are not big, visible adult entomologists of color.*” Community awareness (8.3%) was also prominent; participants repeatedly emphasized that the lack of visible entomologists of color discourages entry into the field. One respondent wrote, “*not seeing someone who looks like you can make you feel like you don’t belong*.”

Structural and systemic barriers comprised 16.7% of coded responses. Participants pointed to socioeconomic inequality (5.6%), structural classism (4.2%), immigration barriers (2.8%), and structural racism (1.4%) as embedded constraints. One respondent articulated this as: “*Housing segregation, imbalanced funding for education, generational wealth… are the biggest hurdles*.” Others referenced visa and passport delays as administrative barriers limiting participation “*probably passport and work visas being approved or held up by government* [sic]*… I hear from my colleagues who recruit that it takes over 6 months to get everything to go through.*”

Identity-based discrimination accounted for 13.9% of responses, with racism alone comprising 8.3% of all coded segments. A respondent described entomology as a field “*dominated by an overwhelming number of white people*,” linking this dominance to bias and stigma. Other forms of discrimination mentioned included sexism, transphobia, and Islamophobia. One respondent summarized these intersecting barriers succinctly: “*Racism, sexism, Islamophobia, transphobia, lack of exposure at K-12*.” This statement was reiterated by the respondent in response to multiple questions.

Outreach-related barriers represented 12.5% of coded responses. Participants emphasized insufficient K–12 exposure, lack of targeted programming, and limited recruitment effort. As one respondent noted, “*It’s hard to recruit students if they have no exposure*.” Another highlighted the lack of institutional recruitment efforts in high schools “*Entomology programs also don’t recruit in high schools etc. (from what I’ve seen), so students may not know that they are an option*.”

Approximately 9.7% of responses reflected social justice avoidance, including color-evasiveness, meritocratic framing, and the assertion that “*no barriers exist*.” For example, one participant reiterated, “*Treat everyone with respect and as an individual and not a representative of a group. Do not patronize others*.” Another stated, “*I do not see any barriers*.”

Finally, one respondent (1.4%) referenced field-specific perceptions, including negative human–insect associations for POC, “*POC are more likely to be from inner city areas where human/arthropod interaction is almost exclusively at a pest level*.”

#### What type of collaboration and mentoring opportunities should be offered to improve retention of Black, Latina/o/x, and Indigenous students?

The most frequently emphasized category for improving retention was Mentorship and Role Models (27%), with participants highlighting the need for role models (12%), mentorship opportunities (12%), faculty guidance (1%), and representation (1%), (Fig 11).

**Fig 11.**
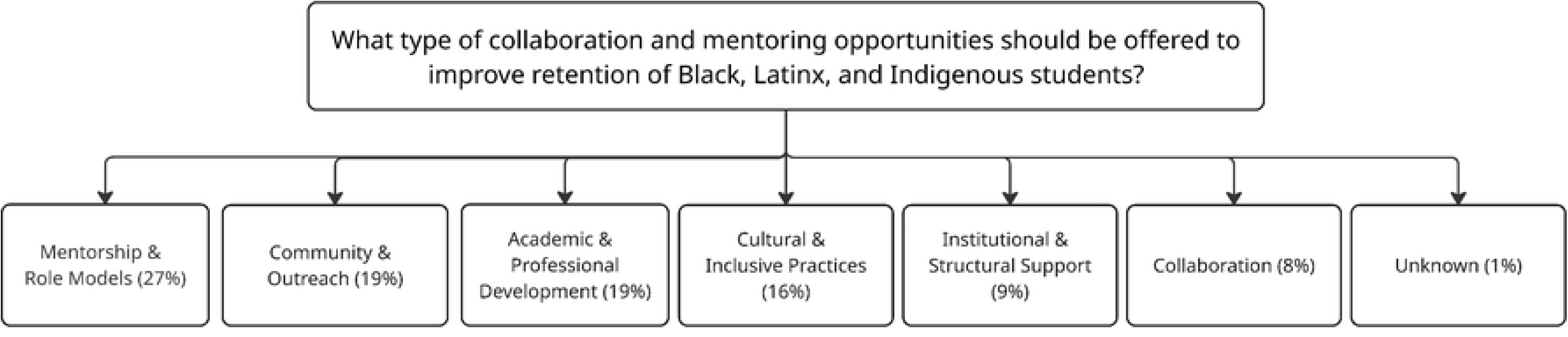
Dendrogram of responses to “What type of collaboration and mentoring opportunities should be offered to improve retention of Black, Latina/o/x, and Indigenous students?” Percentages represent the proportion of coded responses for each theme.

Many respondents stressed the importance of pairing students with successful professionals and supportive faculty. As one participant put it, *“Mentoring is huge! Having a mentor who is willing to listen and learn as well as teach could be an invaluable resource for students.*” Community and Outreach initiatives were also prominent (19%), particularly K–12 outreach (8%) and community engagement (5%) “*You get more buy in if you offer something to the family too,*” along with suggestions for fostering community mindedness, awareness, and cross-sector engagement. Academic and Professional Development emerged as a key area (19%), with particular attention to internship and research experiences (7%), professional networks (4%), resume-building (3%), targeted workshops, youth exposure, and professional development opportunities (each 1%). One respondent stated *“I think it is important to have 1) interdisciplinary collaborations 2) industry or federal agencies on projects for collaborations. This can help widen the scope of opportunities as well as assist with networking for a career after their program is completed.*”

A smaller proportion of responses focused on Institutional and Structural Support (9%), including calls for increased funding (5%) and institutional efforts to remove barriers to access, establish DEI offices, and raise awareness of systemic obstacles (each 1%). For example, “*in college immigration offices which provide mentorship, help, and resources for students and can potentially connect students to professors.*” Cultural and Inclusive Practices accounted for 16% of responses, stressing the importance of cultural awareness (5%), avoiding color evasiveness (4%), and promoting inclusivity, individuality, decolonized curricula, and eliminating tokenization (each 1%). For example, “*Collaborations that lead to publications rather than academic departments and employers just using people of color as diversity tokens.*”

Finally, collaboration was specifically mentioned by 8% of respondents, underscoring the value of cooperative approaches across stakeholders. A small proportion (1%) of responses expressed uncertainty or an inability to identify specific barriers

#### What research areas do you feel should receive more attention in the next 20 years? How has the paradigm of Entomology changed in the past 20 years?

As mentioned in the methods section, these two open-ended questions were analyzed comparatively. Frequency counts were calculated by grouping responses according to the question and thematic category, resulting in a category-by-question summary table (Fig 12).

**Fig 12.**
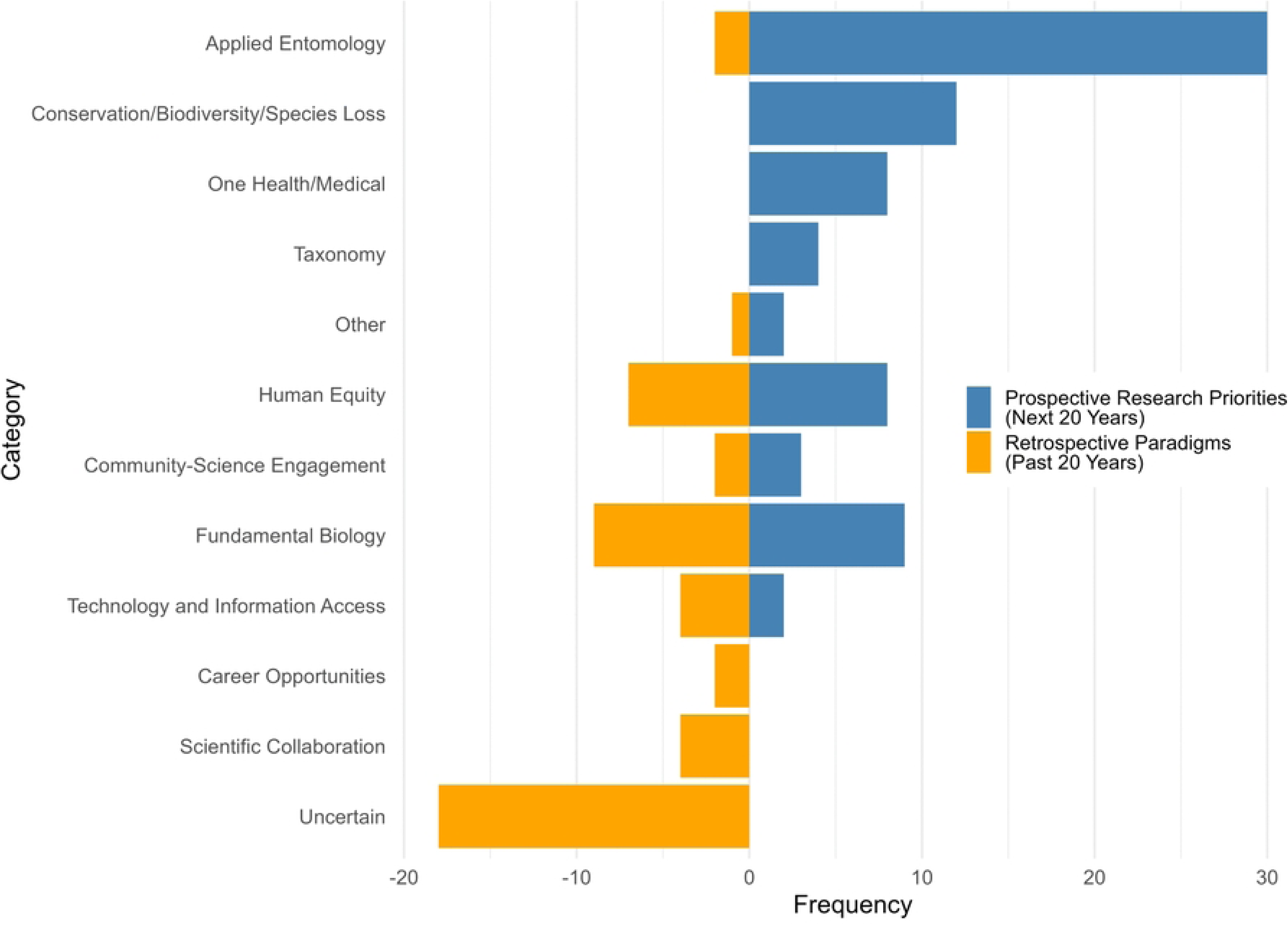
Perceived paradigm shifts and future research priorities in entomology. Bars show the frequency of coded responses across thematic categories comparing changes over the past 20 years with priorities for the next 20 years.

The results show that for the question “*What research areas do you feel should receive more attention in the next 20 years?”,* participants placed the greatest emphasis on Applied Entomology (30 responses, 38.5%) “*Funding for applied research into novel bio-based pesticides*”, followed by concerns about Conservation/Biodiversity/Species Loss (15.4%) and Fundamental Biology (11.5%) “*genomics, phylogenetics*”. Smaller response categories included Human Equity (10.3%) “*ensuring that the field of entomology is accessible to ALL.*” and One Health/Medical (10.3%) “*Medical Urban and Veterinary Entomology and how this research can improve the lives of poor and underserved communities globally*”, while Taxonomy, Community Engagement, Technology and Information Access, and Other appeared less frequently (Fig 12). In contrast, responses to “*How has the paradigm of Entomology changed in the past 20 years*” were dominated by *Uncertainty* (32.7%). It is important to note that the answers in the *Uncertain* category reflect responses such as *“I don’t know, I have only been in the field for a short time*” and “*I’m new to the field,*”. Among more concrete themes, Fundamental Biology (16.4%) *“There is a lot more focus on molecular entomology and genetics, especially in systematics”* and Human Equity (12.7%) *“… more diversity in races of entomologists*” were most common, while Technology and Information Access and Scientific Collaboration (7.3%) also emerged as important. Interestingly, Applied Entomology dropped sharply when considering the future (3.6%) as compared to the past twenty years (38.5%), highlighting a gap between what participants view as priorities for the future and what they perceive as past areas of focus. The category Other, includes codes like Internationalization (retrospective, 1.8%) and Pollination and Funding (prospective, 2.6%).

#### Do you have additional thoughts on the topic of recruitment, training, and retention of Black, Latinx, and Indigenous students into Entomology?

Several participants emphasized the long-standing presence of racism in U.S. institutions, noting that efforts to diversify entomology cannot be fully effective without broader societal change; *“The US as an institution has been racist for its entire existence*.” Others stressed the importance of early exposure to biology and entomology, particularly through K–12 education, to spark interest and encourage future career pathways *“Teaching the younger generation about biology and their surrounding ecosystem may spark their interest in wanting to preserve it*.” The value of non-traditional pathways was also noted, with potential scientists emerging from diverse professional backgrounds, such as culinary or technical fields, where skills like attention to detail and procedural adherence are highly transferable to laboratory settings. Respondents highlighted the role of White allies in advocating for BIPOC colleagues and restructuring power dynamics, while also cautioning that diversity efforts must avoid tokenism or assumptions based solely on race. For example, “*White scientists in positions of power need* [to] *be willing to spend resources on providing better training and recruitment opportunities accessible to BIPOC folks.*” Practical strategies suggested included providing equitable resources based on financial need, improving accessibility and inclusivity within programs (e.g., gender-neutral spaces, DEI-trained staff, professional support), and offering fee waivers for workshops and seminars. For example, *“Financial investment into creating an inclusive environment for when students are involved in programs goes a lot further in retaining students which then leads to their family/friends/community also believing they can succeed in the field*”. Overall, respondents argued that retention should focus on supporting genuinely interested students, fostering an inclusive culture, and creating pathways for individuals from historically marginalized communities to discover and develop their passion for entomology.

## Discussion

This study included students, professors/staff/administrators, industry professionals, retirees, and others. Although the sample is relatively small, the racial/ethnic composition differs meaningfully from both ESA membership and broader national benchmarks. The proportion of Black respondents was notably higher than both ESA’s reported membership demographics and the U.S. population (13). In contrast, Hispanic/Latino/a and Asian respondents were notably underrepresented compared to both ESA membership and the national population. These patterns illustrate not only the difficulty of achieving racially representative sampling in small-scale surveys, but those mismatches also mirror real, long-standing inequalities in how people are represented in science. Targeted oversampling strategies, as used in large national surveys (e.g., NHIS prior to 2016, (30), may offer a practical way forward in social entomology research. By deliberately recruiting from institutions, regions, or networks with higher concentrations of historically excluded groups, studies like this could generate more balanced samples and enable subgroup analyses. Without such strategies, survey efforts risk reproducing the inequities observed not only in professional societies but also in science and engineering education more broadly.

As mentioned before, the results were separated between quantitative and qualitative, the quantitative findings suggest that respondents viewed structural investment as central to advancing inclusion in entomology. The highest mean was observed for Perceptions of Need for Financial and Institutional Support (*M* = 4.85), indicating strong agreement that universities and federal agencies should invest in scholarships and funding for students of color. This category included items such as “*Federal monies should be spent on recruiting Black, Latinx, and Indigenous people into Entomology*” and “*Higher education institutions should create directed scholarship funds towards students of color in STEM/Entomology*”. This concern is consistent with national data showing that economic inequality in the United States continues to vary by race and ethnicity. Hispanic, Black, American Indian and Alaska Native, and Two or More Races populations are overrepresented in the population in poverty (31). NCES further reports that among full-time, full-year undergraduate students, the percentage who received Pell Grants was roughly double for Black and Hispanic students (72% and 82%, respectively), compared with Asian and White students (36% and 34%, respectively) (32). Financial responses differed from those in 2013 which emphasized increased funding but de-emphasized race-conscious decision-making. The response differences between 2013 and 2023 reflects a changing perspective around how race-conscious funding distribution may improve equity.

This theme was followed by Positive Attitudes toward Inclusion (*M* = 3.92), Perceptions of Outreach Strategies & Education (*M* = 3.95), and Perceptions of Need for Early Education *(M*= 3.763), which suggest that respondents view structural investment as essential for broadening participation. This is consistent with national STEM reports emphasizing that broadening participation requires attention to inclusion, access, outreach, mentoring, and support throughout educational pathways. The National Academies (10) noted that increasing participation of underrepresented groups in STEM requires a comprehensive approach spanning preschool through graduate school, rather than isolated interventions alone. Similarly, NSF (33) stated that broadening participation in STEM requires addressing “equity, inclusion, and access in STEM education, training, and careers”. Similarly, in 2013, respondents emphasized early entomology education, mentorship, and public outreach. Together, these findings indicate that mentorship and early exposure remain persistent recruitment needs. Prior literature suggests that these opportunities are shaped by geography: where entomology departments are located influences which communities are most likely to encounter entomologists, outreach programs, and potential mentors (24). Geographic patterns may therefore help explain why early exposure remained a prominent concern in both 2013 and 2023. Given this decade-long consistent response, future entomological outreach programs should focus on providing comprehensive and geographically diverse options for K-12 students.

In 2024, and even in some solicitations published in early 2025, NSF continued to define broadening participation as requiring attention to “equity, inclusion, and access” in STEM (33,34). However, NSF’s updated 2025 (35) priorities reframed these efforts around “merit, competition, equal opportunity, and excellence,” and stated that broadening participation activities must create opportunities for “all Americans everywhere,” while limiting protected-characteristic-based efforts to those explicitly mandated by law. In parallel, Executive Order 14151 directed federal agencies to terminate DEI/DEIA programs and “equity-related grants,” (36) potentially restricting the institutional supports and scholarships that respondents in this study identified as essential for recruitment and retention. Reports have documented the cancellation or termination of NSF grants that included DEI-focused or broadening participation components in STEM (37–40). These developments create tension between the strong support for structural investment observed in this sample and the broader national context that shapes the availability of resources to implement such efforts.

Awareness of Role Models & Diversity Resources (*M* = 2.45), indicated a persistent visibility gap where people value diversity but may not know the historical or contemporary figures who embody it. This indicates limited familiarity with platforms such as Entomologists of Color and with historically significant figures like Dr. Charles Henry Turner. Visibility gaps are consistent with longstanding concerns about the lack of same-race mentors and the underrepresentation of BIPOC faculty in biological sciences (41,42). Notably, similar issues were identified in the 2013 study by Abramson et al. in which respondents emphasized the need for role models and early exposure to entomology. Persistent lack of awareness a decade later suggests that the field has not effectively promoted its diverse history or expanded access to culturally relevant mentorship. Such gaps may hinder belonging and contribute to feelings of isolation among underrepresented groups, even as contemporary respondents express strong support for institutional inclusion efforts.

Group differences were most pronounced for Perceived Workplace Discrimination, which varied by race/ethnicity: Black or African American participants reported substantially higher levels of perceived workplace discrimination. This pattern aligns with extensive literature documenting racialized experiences in STEM, where BIPOC students and professionals encounter exclusionary climates, limited representation, and systemic barriers (3,6,7). Similarly, a gender effect emerged for Perceived Workplace Discrimination, with women reporting higher discrimination than men. However, non-binary/third gender responses did not differ significantly, likely due to the small respondent population. In 2013, respondents shared their experiences of discrimination as well, demonstrating discrimination as a consistent barrier to equity.

The qualitative findings help contextualize the quantitative results by demonstrating that respondents understood inclusion in entomology is shaped not only by recruitment, but also by the conditions that sustain participation over time. Across responses, participants identified DEI as the most pressing future challenge to advancing equity in entomology and repeatedly pointed to structural and systemic barriers rather than individual-level challenges. Limited mentorship guidance, and visible role models stood out as the strongest recurring themes across questions. Respondents repeatedly stressed supportive mentors, representation, and faculty guidance as essential for persistence in the discipline.

This aligns with prior work suggesting that disparities in STEM fields may emerge less from initial interest than from unequal retention and support. For example, Beltran et al. (1) investigated the role of field course participation in shaping academic outcomes and self-efficacy among underrepresented students in EEB. Using data from more than 29,000 undergraduates at UC Santa Cruz (2008–2019), they found no demographic differences in intended majors at admission. However, five years later, underrepresented students were significantly less likely to graduate with EEB degrees, suggesting that inequities in matriculation rates are driven more by retention than recruitment. One contributing factor is that participation in field courses was lower among HEU groups, yet such courses were linked to increased self-efficacy, stronger retention in EEB majors, higher likelihood of college graduation, and improved GPAs.

In this context, respondents’ emphasis on mentorship, outreach, funding, and institutional support suggests an understanding that broadening participation requires more than attracting students into the field; it also requires creating the material and social conditions that make persistence possible. This seems to be a cultural shift between 2013 and 2023; in 2013, respondents focused on recruitment (e.g., outreach, K-12 activities, increasing awareness) whereas in 2023 respondents addressed these issues while providing situative context that recruitment alone is not sufficient (e.g., funding, employment, mentorship, role models, retention). This may be a product of both the changing cultural context around DEI in the U.S. and the reenvisioned survey which centered both structural and individual-level concerns.

The qualitative data also suggests that respondents viewed culture and climate as central to retention. In particular, concerns about tokenism, lack of belonging, and the need to challenge color-evasive “meritocracy” rhetoric suggest that exclusion in entomology is reproduced not only through resource inequities, but also through everyday institutional norms. Color-evasiveness has been described as a contemporary form of racism that obscures structural inequality while resisting race-conscious change (8,9). Applied to entomology, such color evasion may weaken access to culturally relevant mentorship, reduce opportunities for meaningful belonging, and make inequities appear as individual rather than institutional problems. Respondents called for both structural investment and cultural change, suggesting that participants viewed recruitment, retention, and inclusion as inseparable from the broader climate in which training occurs.

Finally, respondents prioritized applied entomology for the next 20 years—along with conservation/biodiversity and fundamental biology. However, when respondents were asked to reflect on the past 20 years, many respondents were uncertain. Among those who answered more concretely, they most often mentioned fundamental biology and human equity. This suggests a gap between where respondents think the field should go and how they understand its recent trajectory.

### Comparisons with the 2013 study

This study was conceived from the beginning not merely as a replication of the Abramson et al. (25) study, but as a temporal comparison of evolving paradigms in entomology. Its purpose was to examine how perceptions of recruitment, training, retention, and inclusion have developed over the last decade. The 2010’s and early 2020’s were characterized by important shifts in STEM and diversity, equity, and inclusion (DEI) discourse including increased attention to systemic racism, institutional climates, and broad social movements and efforts to increase equity in society.

During this period, entomology also faced growing concerns regarding declining enrollment (14), climate change (6), insect biodiversity decline (43,44), and workforce sustainability (24), making it particularly important to reassess how the field conceptualizes recruitment and inclusion.

Accordingly, the comparison is valuable not only because it reveals continuity in longstanding concerns, but also because it highlights a conceptual shift in how those concerns are understood (i.e., intrinsic vs. extrinsic factors to underrepresentation). Perceptions of underrepresentation in entomology have moved from a focus on individual preparation and participation toward broader recognition that institutional climate, systemic inequities, and disciplinary norms shape who enters, persists, and advances in the field. This interpretation parallels broader STEM literature, where underrepresentation is increasingly understood not only as a “pipeline” problem but also as a consequence of institutional and cultural conditions (1,3,6,7).

The Abramson et al. (25) study focused on Black and Hispanic entomologists and reported item-level means across recruitment and training domains, whereas the 2023 study broadened racial/ethnic representation and participant roles (students, faculty/staff, industry, retirees). While some comparisons are limited by the original study’s sample size and methodology, the comparison remains valuable for understanding how perceptions evolved between 2013 and 2023.

Notably, mentorship was identified by 2013 respondents as the single most influential factor, alongside precollege interventions such as tutoring, outreach programs, and increased visibility of entomology in secondary education. These priorities remained highly salient in the 2023 findings, where K–12 outreach emerged as the most frequently cited recruitment strategy (Fig 9) and mentorship and role models constituted the most prominent theme for retention (Fig 11). However, K-12 outreach cannot exist in isolation without structural change toward equitable educational environments.

The comparison also reveals persistent tensions that have remained largely unresolved. In the 2013 study, respondents expressed a paradoxical stance stating that the “majority of respondents strongly agreed that efforts should be made to recruit Black and Hispanic students into entomology but the respondents were significantly less aligned on the methods”… “they did not agree, however, whether ethnicity should be a factor in recruiting” (25). This seemingly contradictory perception was echoed in the 2023 data through the continued presence of color-evasive and meritocratic responses although these occurred less often than structurally oriented explanations. For example, color-evasive and meritocratic themes accounted for 5.0% of coded segments in responses about recruitment strategies and were present within 9.7% of coded segments addressing barriers to recruitment (Fig 9 and 10). This contradiction suggests that ideological divisions regarding equity-focused interventions persist within the field and even within individuals. While awareness of structural inequities has increased since 2013, consensus on how to address them remains incomplete, reflecting broader debates within STEM regarding the role of targeted versus universal approaches to equity.

Beyond recruitment and inclusion, the comparison also suggests a shift in how respondents understood the future of entomology. In Abramson et al. (25), concerns were primarily discipline-centered, focusing on funding, the decline of taxonomy and systematics, loss of entomology programs, and applied issues such as pest management, insect resistance, and invasive species. By 2023, these concerns had broadened to include not only funding and workforce sustainability, but also ecological, institutional, and cultural challenges. Respondents highlighted climate change, insect decline, disciplinary fragmentation, weak quantitative preparation, poor labor conditions, and mental health as important issues. Most notably, diversity, equity, and inclusion emerged as the largest area of concern in 2023, with explicit references to racism, unwelcoming environments, and systemic barriers (Fig 7). Together, these findings suggest that perceptions of the field have shifted from a primarily disciplinary focus toward a more integrated view of entomology as shaped by broader environmental, institutional, and social forces.

Finally, the comparison of research priorities and perceived paradigm shifts between 2013 and 2023 further illustrates both continuity and change in the field. In Abramson et al. (25), respondents emphasized applied and discipline-specific areas, including integrated pest management, insect–plant interactions, taxonomy and systematics, and vector control, with emerging attention to molecular techniques and global environmental issues such as climate change. Similarly, the 2023 findings indicate that applied entomology remains a dominant priority, alongside interest in conservation, biodiversity, fundamental biology, human equity and One Health, reflecting a broader integration of social and global health perspectives into entomological research since 2013 (Fig 12).

The 2023 findings suggest that the future of entomology may depend on the field’s ability to integrate scientific innovation with institutional and cultural change. Respondents framed future challenges not only in terms of technical expertise, but also through inclusivity, workplace climate, interdisciplinary collaboration, mental health, and ecological change. This suggests that recruitment and retention efforts must address both scientific priorities and the conditions needed to sustain a diverse entomological workforce.

Taken together, these findings suggest that the decade between 2013 and 2023 represents not a simple progression but a complex transformation in how diversity and the future of entomology are understood. This shift has important implications for the future of the field, indicating that efforts to broaden participation must include sustained attention to climate, equity, and systemic reform. In this sense, the 2023 study not only confirms the enduring importance of established strategies but also highlights the necessity of integrating these approaches within a broader, more critical understanding of the barriers shaping participation in entomology.

## Implications and Conclusion

### Strategies for early exposure to science

The findings of this study have important implications for efforts to broaden participation in entomology and STEM more broadly. Consistent with prior work, early exposure and mentorship remain central mechanisms for recruitment and retention. The persistence of disparities in participation suggests a need for a stronger emphasis on retention, institutional climate, and long-term support systems. Participants identified structural barriers-including racism, unwelcoming environments, funding inequities, and reliance on unpaid or undercompensated labor-as critical challenges, indicating that broadening participation requires systemic, rather than solely individual-level, solutions.

These findings further highlight the need for institutional accountability in addressing inequities within entomology. Mentorship, training and research must be grounded in institutionalized and formalized anti-racist policies (45). As Kendi, (46) argues, anti-racism is an ongoing process that requires continuous reflection and learning. Meaningful progress depends on humility, a willingness to learn from mistakes, and sustained engagement at both individual and organizational levels.

Importantly, the continued presence of color-evasive and meritocratic perspectives in the data suggests that ideological tensions regarding equity-focused interventions persist, underscoring the need for ongoing dialogue and education within the field. At the same time, concerns related to compensation, workload, and early-career instability indicate that academic labor conditions represent a significant, yet often overlooked, barrier to retention, particularly for individuals from historically marginalized backgrounds.

Finally, the evolving research priorities identified in this study point to a broader transformation in the scope and relevance of entomology. The increasing emphasis on global challenges such as climate change, biodiversity loss, One Health, and human equity suggests that the future of the discipline will depend on its ability to engage with interdisciplinary and socially relevant research agendas. These findings suggest that advancing diversity in entomology will require not only sustained investment in mentorship and early exposure, but also structural reform, inclusive training environments, and alignment with broader societal and environmental priorities.

### Limitations and future directions

Interpretation of these findings is tempered by some limitations. The small, non-probability sample and uneven subgroup sizes limit statistical power for post-hoc contrasts and constrain generalizability. However, the uneven subgroup sizes closely mirrored ESA and/or US population demographics with some exceptions as described in the Methods. The surveys offered in 2013 and 2023 were similar but not identical and methods varied between the two studies which limit comparative conclusions. Participants in the 2023 study may have also been primed to think about DEI topics by the consent form which provided context about the study and its aims. As in any survey, some of the questions may not have reflected the specific context or environment of which respondents were familiar. For example, few respondents were familiar with Dr. Charles Henry Turner but may have been familiar with other entomologists of color that were not explicitly listed. Future research should employ targeted oversampling and stratified recruitment to balance samples as much as possible. Entomology is a relatively small and specialized disciplinary field that relies on recruitment from other disciplines. Future studies may aim to understand how entomology is distinct from related fields regarding equity, recruitment, and retention.

## Supporting information

**S1 Table. Scale assessing perceptions and attitudes toward inclusion in entomology.** This table lists the items included in each subscale.

**S1 Fig. Complete dendrogram of respondents’ perceived significant challenges for the future of entomology.** Full hierarchical coding structure corresponding to the condensed dendrogram presented in Fig 7.

**S2 Fig. Complete dendrogram of respondents’ perceived significant challenges in training young Black, Latinx, and Indigenous students.** Full hierarchical coding structure corresponding to the condensed dendrogram presented in Fig 8.

**S3 Fig. Complete dendrogram of strategies to recruit Black, Latinx, and Indigenous students into entomology.** Full hierarchical coding structure corresponding to the condensed dendrogram presented in Fig 9.

**S4 Fig. Complete dendrogram of barriers to recruitment of Black, Latinx, and Indigenous students into entomology.** Full hierarchical coding structure corresponding to the condensed dendrogram presented in Fig 10.

**S5 Fig. Complete dendrogram of collaboration and mentoring opportunities to improve retention of Black, Latinx, and Indigenous students.** Full hierarchical coding structure corresponding to the condensed dendrogram presented in Fig 11.

